# Targeting NF-κB prevents Schwann cell reprogramming and plexiform neurofibroma initiation in mouse model of Nf1

**DOI:** 10.64898/2026.11.04.764357

**Authors:** Ramya Ravindran, Eui-Kyung Youn, Noemi Kedei, Kwangmin Choi, Avery Volz, Eva Dombi, Jay Pundavela, MiOk Kim, Robert J. Spinner, Sara Szabo, David A. Largaespada, Jack F. Shern, Nancy Ratner

## Abstract

Plexiform neurofibromas arise in approximately half of all individuals with Neurofibromatosis Type I (NF1). Genetic lesions apart from *NF1* pathogenic variants are not present in *NF1* mutant Schwann cells, suggesting that tumorigenesis may instead require cellular rewiring. To identify putatively re-wired tumor cells, we used multiplex imaging and flow cytometry to delineate markers that distinguish tumor Schwann cell from non-tumor Schwann cells. We identified tumor Schwann cells as CD44+;CD49f+ cells that show increased transcription of epithelial to mesenchymal plasticity (EMP)-associated genes. CD44+;CD49f+ expression was induced by treating *Nf1-/-*Schwann cells with stressors or by direct NF-κB pathway activation via IKK2; treated Schwann cells increased secretion of NF-κB targets and immune cell chemoattractants CCL2 and CCL3. In established tumors, MEK inhibition plus NF-κB blockade diminished tumor cell proliferation, tumor Schwann cells, immune cell populations, tumor cytokines and tumor number. Notably, transient single agent NF-κB pathway blockade was sufficient to significantly delay tumor formation in mice. Thus, *Nf1* loss in Schwann cells is followed by NF-κB activation and cellular reprogramming that drives tumor initiation and targeting NF-κB activation can reduce NF1-deficient tumor formation.

## Introduction

Neurofibromatosis type I (NF1) is one of the most common tumor predisposition syndromes in humans, affecting approximately 1 in 3000 individuals worldwide (1). Biallelic loss-of-function mutations in the *NF1* gene in Schwann cell lineage cells are present in neurofibromas – benign tumors that develop in the nerves of the peripheral nervous system (2). Plexiform neurofibromas (PN) associated with nerve trunks arise in individuals with NF1, can cause significant morbidity through nerve compression, disfigurement and pain, and can progress to an aggressive malignancy known as malignant peripheral nerve sheath tumor (MPNST) (3,4). Despite this clinical burden, therapeutic options are limited. The *NF1* gene product, neurofibromin (NF1), functions as a GTPase-activating protein that accelerates the hydrolysis of active RAS-GTP into inactive RAS-GDP, thereby attenuating RAS signaling (5). The loss of NF1 results in elevated intracellular RAS-GTP levels (6–8) and increased signaling through down-stream effector pathways, particularly the mitogen-activated protein kinase (MAPK) cascade (9). Indeed, MEK inhibition, which targets the activated Ras/MAPK pathway downstream of NF1 loss, is the only FDA approved pharmacological treatment for PN. MEK inhibitor therapy only achieves partial tumor volume shrinkage and requires sustained administration to prevent regrowth (10–13). The inability to achieve durable tumor shrinkage or to prevent tumor formation underscores a need to understand the earliest molecular events that drive PN initiation.

Mouse models have been developed that are excellent mimics of the histology and growth kinetics of human neurofibromas (14–20). These genetically engineered mouse models (GEMM) have been instrumental in establishing that biallelic *Nf1* loss in Schwann cells is necessary for PN formation and, because they are autochthonous lesions, provide opportunities to study how neurofibromas form. In the *DhhCre;Nf1fl/fl* mouse model, Cre-recombinase-mediated loss of *Nf1* occurs in Dhh-expressing Schwann cell precursor cells mid-gestation, yet tumors do not form until 4 months of age (18). This latency cannot be explained by the acquisition of additional genetic mutations - beyond biallelic mutation/loss of *NF1*, no additional genetic lesions have been identified in PN Schwann cells (21–25). The finding that *NF1* loss in Schwann cells is necessary but not sufficient for PN initiation implies that non-mutational, cell-intrinsic or microenvironmental reprogramming events must occur in the interval between *Nf1* loss and tumor onset. Identifying such permissive events is central to understanding PN biology and is predicted to identify strategies that could prevent tumor formation in at-risk individuals.

The tumor microenvironment plays an essential and active role in PN formation. *Nf1-/-*Schwann cells secrete elevated levels of growth factors and cytokines including SCF, CCL2, and CXCL10, factors that recruit mast cells, macrophages, dendritic cells and T cells (26–29). Each of these cell types is present in PN; immune cells comprise about 30% of PN cells (30–34). These immune cells contribute to tumor development. For example, in a mouse model, the absence of T cells prevents PN formation, and tumors form after replacement of CD8 T cells, demonstrating that this component of the adaptive immune system is an active driver of tumorigenesis (29,35). Fibroblasts are also abundant in neurofibromas, constituting 20-30% of neurofibroma cells, and neurofibromas contain a tumor matrix rich in collagens and laminins (14,17,36,37). Signaling pathways active in *Nf1-/-*tumor Schwann cells that initiate and orchestrate stromal and immune remodeling, particularly prior to tumor formation, remain unknown.

A candidate underexplored signaling pathway in PN tumorigenesis is nuclear factor-κB (NF-κB) signaling. NF-κB transcription factors are master regulators of inflammation, cell survival, and stress responses, which are increasingly recognized as drivers of tumor initiation through their capacity to reprogram epithelial and stromal cells, induce expression of pro-tumorigenic cytokines and initiate epithelial to mesenchymal plasticity (EMP) properties (38,39). In the canonical pathway, extracellular signaling including cytokines, growth factors, and oncogenic stress activate the IKK complex which phosphorylates IkBa, targeting it for proteasomal degradation and releasing p65/RelA for nuclear translocation and target gene activation (38,40). In peripheral nerve Schwann cells, NF-κB has been shown to regulate inflammatory chemokine production following nerve injury, and IKK deletion in non-myelinating Schwann cells reduced CCL2 and CXCL10 expression (41,42) –chemokines that contribute to immune cell recruitment in PN (29,43,44). Importantly, a recent transcriptomic study comparing *Nf1* mutant Schwann cells before and after tumor formation predicted NF-κB pathway upregulation in tumor Schwann cells and detected nuclear p65 protein in mouse and human PN tissue sections (34). However, whether NF-κB is a cause or consequence of PN formation, how and whether it might work together with Ras pathway activation in tumorigenesis, and whether it represents a therapeutically actionable step in tumorigenesis have not been studied.

Here we demonstrate that NF-κB activation is a required and targetable initiating event in PN formation. We identify CD44 and CD49f as surface markers enriched in transcriptionally reprogrammed tumor Schwann cells. We show that this reprogrammed state can be induced by NF-κB activation in *Nf1-/-*Schwann cells, and that it occurs concurrently with nuclear p65 accumulation at the onset of tumor formation in vivo, preceded by an increase in collagen deposition and followed by infiltration of immune cells and an expansion of cancer-associated fibroblasts. Remarkably, transient blockade of NF-κB signaling prior to tumor onset significantly delayed plexiform neurofibroma initiation – establishing NF-κB as a rate limiting step in PN formation. Targeting this pathway suggests a new therapeutic paradigm focused on preventing tumor initiation rather than reversing established disease.

## Results

### Multiplex single-cell spatial imaging visualizes diverse cell types in plexiform neurofibroma and identifies CD44+;CD49f+ tumor cells

To identify immune and non-immune cell types in plexiform neurofibroma, we spatially analyzed *DhhCre;Nf1fl/fl* mouse PNs at single cell resolution using the Phenocycler [formerly CODEX (45)] multiplex imaging system (Figure 1A). We obtained unfixed frozen tissue sections from 14 PNs, collected from 8 mice. Samples were stained with a 27-plex antibody panel optimized for use in PN tissue; Supplemental Table 1 shows the panel and the order of antibody application in sequential cycles with appropriate fluorophores. In the *DhhCre;Nf1fl/fl* mouse model PNs form within nerve regions adjacent to some, but not all, dorsal root ganglia (DRG) (18). These sensory ganglia contain neuronal cell bodies and associated satellite cells (a type of Schwann cell), while nerves contain Schwann cells aligned upon nerve axons. DRG and nerve are each surrounded by a barrier called the perineurium, made up of fibroblasts that express the marker CD49f/ITGA6. In *DhhCre;Nf1fl/fl* animals all nerves are enlarged (hyperplastic) (18). To analyze cell composition in tumors, we separately marked DRG, nerve, and tumor regions of interest (ROI) for cell type identification (Figure 1B), then focused on tumor ROIs (n=14). Tumor ROIs were segmented using Hoechst dye as a nuclear marker, and expanding to define cell surface and segment each cell (total of 156,074 cells were segmented). Identification of the ROIs was based on consultation with pathologist, Dr. Sara Szabo, MD, and based on hematoxylin-eosin (H&E) analysis of adjacent tissue sections, using established criteria (46).

**Figure 1.**
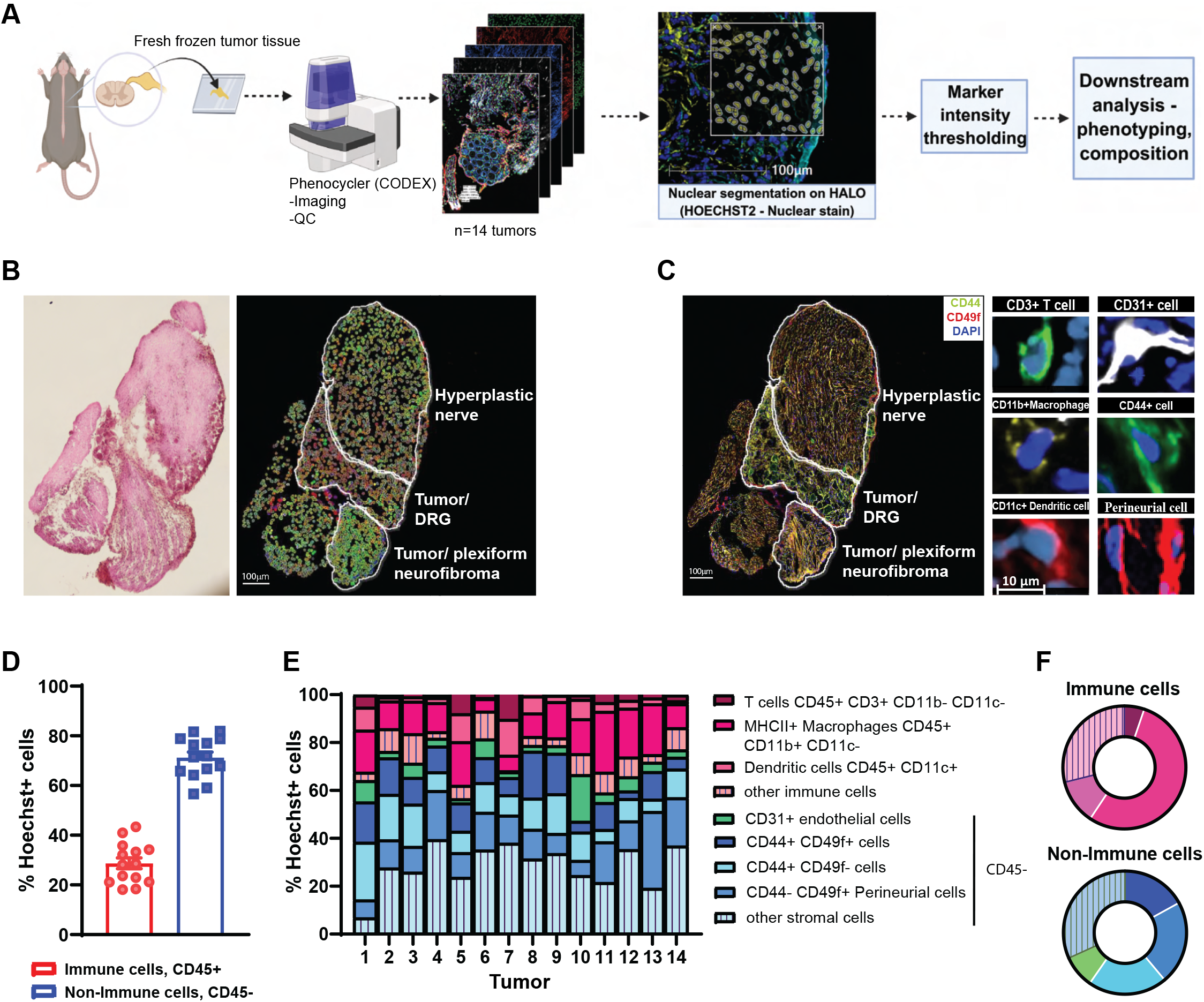
Visualization of tumor cell types in plexiform neurofibroma using single-cell spatial imaging. (A) Schematic overview of the workflow (Created with BioRender.com) (B) Hematoxylin-eosin (H&E) staining (left) showing the tumor, nerve and adjacent Dorsal Root Ganglion (DRG) in tissue and representative cell masks (left) of identified cell types based on surface marker co-expression (right) with ROIs indicating analysis regions. Scale bars, 100μm. (C) Representative Phenocycler stained tumor tissue (left) and single cell images of identified cell types (right). Scale bars are indicated in μm. (D) Bar plot of percent immune and non-immune cell populations in tumor (*n* = 14). (E) Stacked bar plots per sample clustered by percentage of each identified subpopulation in the neurofibroma environment. (F) Doughnut plot showing proportions of the immune and non-immune subpopulations in neurofibroma based on marker analysis. See also Supplemental Table 1.

Individual cells within tumor ROIs were next identified based on marker co-expression (Figure 1C). Immune cells were defined based on expression of CD45; Figure 1D shows that the proportion of CD45+ immune and CD45-non-immune cells in each of the analyzed tumor ROI is similar. CD45+ immune cells were categorized as CD3+ T cells, CD11b+;MHCII+ macrophages and CD11c+ dendritic cells in each of the 14 analyzed ROIs (Figure 1E). The non-immune (CD45-) cells in PN include Schwann cells, fibroblasts and endothelial cells (Figure 1, E-F). CD31 identified endothelial cells. We then further analyzed the non-immune cell subpopulations that clustered as CD44-;CD49f+ as perineurial cells; CD49f identifies perineurial cells in normal nerve (47). Unexpectedly, this spatial analysis also identified large numbers of CD44+;CD49f-cells and CD44+;CD49f+ cells in tumor. CD44 can be expressed by inflammatory cells and fibroblasts and is often upregulated at sites of injury (48,49). Indeed, Schwann cells express CD44 during early development and after *Nf1* loss (50). Given that CD44+;CD49f+ cells expressed the Schwann cell/fibroblast marker CD44 and the perineurial cell marker CD49f we sought to better define these non-immune PN cell populations, and test if they are tumor specific.

### Schwann cells in PN uniquely co-express the cell surface markers CD44 and CD49f

To better characterize the non-immune cell subpopulations defined by spatial analysis in plexiform neurofibroma we developed a flow cytometry approach (Supplemental Figure 1A). Single cells were generated from visible tumors, and from adjacent DRG/nerve that had not formed tumors (designated non-tumor DRG) from the same *DhhCre;Nf1fl/fl* mice. We also generated single cells from wildtype mice DRG/nerve. Consistent with the spatial analysis in Figure 1, CD44+;CD49f+ cells made up 20% of tumor non-immune cells but were rare in wild type and non-tumor DRG samples (Figure 2, A-B). To determine if these tumor-enriched CD44+;CD49f+ cells are Schwann cells we carried out cell-type marker analysis. DhhCre expressing (“Lineage+”) cells that undergo Cre-recombinase-mediated loss of *Nf1* were identified by expression of the GFP reporter in nerve Schwann cells and endoneurial, but not perineurial, fibroblasts (51). Of total live (immune + non-immune) tumor cells, about 7% expressed this lineage marker (Figure 2C and Supplemental Figure 1B). Most lineage+ cells (80%) were CD44+;CD49f+, suggesting that they are Schwann cells. To confirm this, we analyzed expression of GFAP. In mouse DRG/nerve, GFAP expression is limited to non-myelinating Schwann cells and some satellite glial cells. Most GFAP+ cells in tumors were CD44+;CD49f+ (Figure 2D). Of GFAP+ and lineage+ cells 80% were CD44+;CD49f+ cells, confirming that most or all are Schwann cell-lineage cells. p75/NGFR expression is often used as an additional label for non-myelinating Schwann cells; 12% of live tumor cells were p75/NGFR+ (Figure 2C and Supplemental Figure 1B); of p75/NGFR+ cells a third expressed CD44+;CD49f+ (Figure 2D).

**Figure 2.**
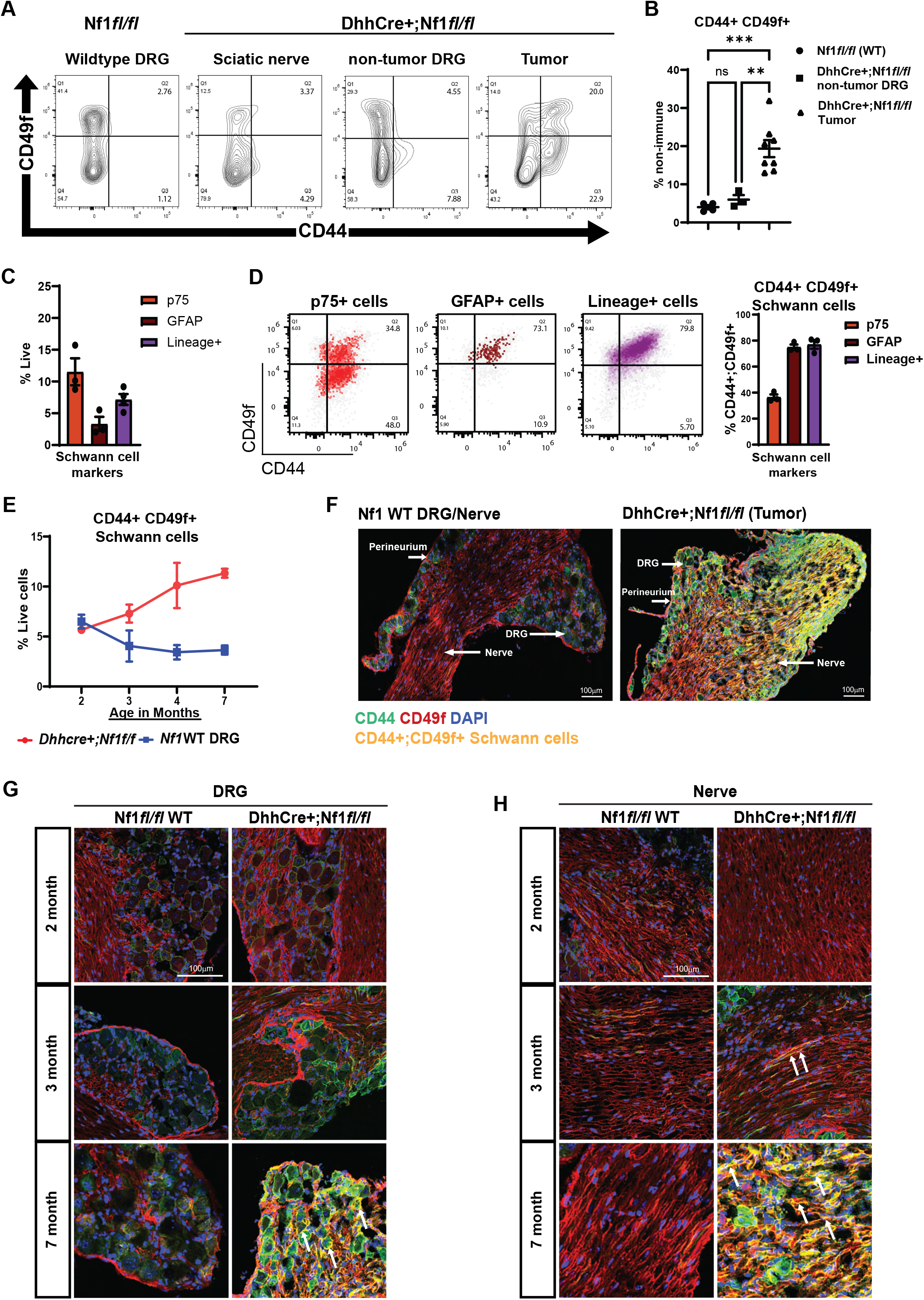
Schwann cells co-express CD44 and CD49f surface markers in tumor. (A) Contour plots from representative flow cytometry samples showing the non-immune CD44+ CD49f+ cell population as increased only in the tumor. (B) Scatter plot showing the CD44+CD49f+ cell population as a percent of the non-immune cells. Tukey’s test corrected *p* values were determined using Ordinary One-Way ANOVA. \**p*<0.0332, \*\**p*<0.0021, \*\*\**p*<0.0002, \*\*\*\**p*<0.0001, ns [non-significant]. (C) Bar plot showing proportions of p75, GFAP and Dhh lineage (Schwann cell markers) positive cells as a percent of Live cells as quantified by flow cytometry. (D) Density dot plots of Schwann cell populations showing co-expression of CD44 and CD49f in a representative sample. Right panel shows bar plot of the Schwann cells that coexpress CD44 and CD49f. (E) Flow cytometry analysis of CD44+ CD49f+ Schwann cells (*p*<0.01 for 4- and 7-months) in *DhhCre+;Nf1fl/fl* mouse DRG/ tumor at ages 2, 3, 4(tumor onset) and 7 months (n = 2-4 per group). Tukey’s test corrected *p* values were determined by Ordinary Two-Way ANOVA. (F) Immunofluoresence staining for CD44+; CD49f+ Schwann cells in *DhhCre+;Nf1fl/fl* plexiform neurofibroma vs Nf1 WT DRG. Scale bars, 100μm. (G) Representative images of tumor DRG regions and (H) tumor nerve regions at ages 2-3- and 7-months showing CD44+; CD49f+. Scale bars, 100μm. See also Supplemental Figure 1.

To test if CD44+;CD49f+ cells are present before tumor formation, we analyzed cells by flow cytometry in mice at 2, 3, 4, and 7 months of age (Figure 2E). Of note, tumor formation in this mouse model is visible at 4 months of age (18). Double labelled cells constituted a small population of cells in wild type samples, which declined at later time points. In *DhhCre;Nf1fl/fl* samples CD44+;CD49f+ cells increased concurrent with the onset of tumorigenesis, in 4-month-old mutant animals. We next stained tissue sections from age-matched wildtype controls (*Nf1fl/fl*) and tumor bearing *DhhCre;Nf1fl/fl* mice. Tumors, but not wildtype nerve/DRG, contained significant numbers of double positive cells (Figure 2F). Further analysis in tissue sections from mice at 2, 3, or 7 months showed that CD44+;CD49f+ expression is abundant in the DRG in satellite glial cells and in tumor cells at the 7-months time point. Rarer double-positive cells were detected in pre-tumor nerve at the 2 and 3-month time points (Figure 2, G-H).

We wondered if a CD44+;CD49f+ Schwann cell population is also present in human plexiform neurofibroma. To test this, we analyzed dissociated single tumor cells from 4 patient samples, stained for Schwann cells (p75/NGFR), fibroblasts (CD34) and immune cells (CD45). Schwann cells (p75/NGFR+) accounted for 25% of PN cells; fibroblasts (CD34+) constituted 24% (Figure 3A) and CD45+ immune cells for 40% of the total live cells. We next test if CD44+ and CD49f+ mark these populations. CD44+;CD49f+ cells made up 12% of total live tumor cells (Figure 3B). Of the CD44+;CD49f+ cells in human PN, 80% were p75+, confirming their Schwann cell identity (Figure 3C) and of these, surprisingly, 80% also expressed the mesenchymal marker CD34 (Figure 3D). Together, these spatial and flow cytometry analyses show that CD44+;CD49f+ cells are reprogrammed tumor Schwann cells, express mesenchymal markers, and increase in neurofibromas.

**Figure 3.**
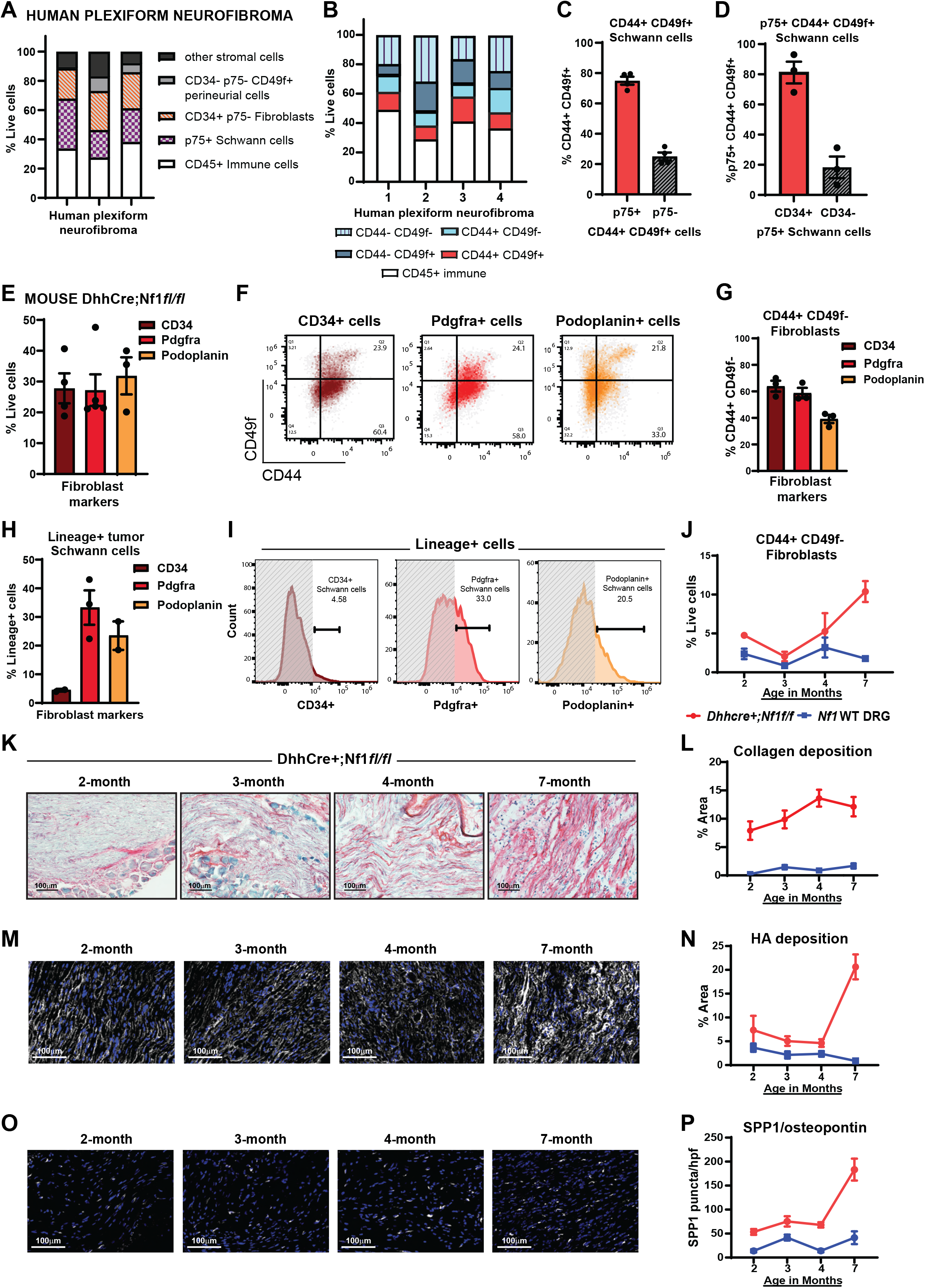
Plexiform neurofibromas increase fibroblast cell populations after tumor onset and Schwann cell EMT. (A) Stacked bar plots of total Schwann cells, fibroblasts, immune cells and other stromal cells in human plexiform neurofibroma as identified by flow cytometry analysis (n=3 samples). (B) Stacked bar plots per sample clustered by percentage of each identified subpopulation in human plexiform neurofibroma by flow cytometry analysis (n=4 samples). (C) Quantification of p75+ Schwann cells by flow cytometry in the CD44+;CD49f+ human plexiform neurofibroma sub-population (n=4 samples). (D) Percent quantification of CD34+ mesenchymal marker expression on p75+;CD44+;CD49f+ Schwann cells in human plexiform neurofibroma (n=3 samples). (E) Proportions of CD34, Pdgfra and Podoplanin (Fibroblast markers) positive cells as a percent of the Live cell population in *DhhCre+;Nf1fl/fl* mouse plexiform neurofibromas. (F) Representative density dot plots of CD44+CD49f-Fibroblast cell populations. Bar plot (G) quantifies the fibroblasts that are CD44+ CD49f-. (H) Quantification of mesenchymal marker (CD34, Pdgfra, Podoplanin) expression on Lineage+ Schwann cells, and representative histograms (I) showing marker expression on the Schwann cells. (J) Flow cytometry analysis of CD44+ CD49f-Fibroblasts (*p*<0.0001 for 7-months) in *DhhCre+;Nf1fl/fl* mouse DRG/ tumor at ages 2, 3, 4 (tumor onset) and 7 months (n=2-4 per group). (K) Sirius red staining showing changes in collagen deposition (red) at ages 2-7 months in *DhhCre+;Nf1fl/fl* mice. Scale bars, 100μm. Average collagen deposition per unit area in (L) (n=3 per group). *p*<0.0005 for all comparisons. (M) Representative images of hyaluronic acid (HA) deposition at ages 2-7 months (*p*<0.0001 for 7-months) in the *DhhCre+;Nf1fl/fl* mouse model as identified by HA binding protein (HABP) immunostaining. Scale bars, 100μm. Average HA deposition is quantifies per unit area in (N) (n=3 per group). (O) SPP1/ osteopontin staining at ages 2-7 months (*p*<0.0021 for 4-months, *p*<0.0001 for 7-months) in *DhhCre+;Nf1fl/fl* mice. Scale bars, 100μm. Staining quantified as puncta per high power field (hpf) as SPP1 is both intra- and extracellular (P) (n=3 per group). Significance determined with Ordinary two-way ANOVA and Tukey’s multiple comparisons test. See also Supplemental Figure 1.

### Schwann cell reprogramming and collagen deposition precede CD44+;CD49f-CAF expansion in PNs

In human p75/NGFR is an accepted Schwann cell marker. However, in mouse tumors, many p75/NGFR+ tumor cells were CD44+ but did not express CD49f, GFAP, or the lineage marker (48%, Figure 2D). To test if these cells are fibroblasts we stained with the fibroblast markers CD34 and PDGFRα; both labeled the CD44+;CD49f-cells (Figure 3, E-G and Supplemental Figure 1C). We verified this further using podoplanin, which marks tumor-associated fibroblasts (52), defining the CD44+;CD49f-mouse neurofibroma cells as fibroblasts. Surprisingly, 20% of the lineage+; GFAP+; CD44+;CD49f+ tumor Schwann cells also expressed podoplanin, and 30% expressed PDGFRα (Figure 3, H-I). Thus, consistent with the human PN flow cytometry data, mouse tumor Schwann cells aberrantly express mesenchymal markers. We use the term epithelial-mesenchymal plasticity “EMP” to classify the Schwann cell reprogrammed state we observe in PN, a partial epithelial-mesenchymal transformation (EMT) (53–55).

CD44+;CD49f-(podoplanin+;PDGFRα+;CD34+) fibroblasts increased in *DhhCre;Nf1fl/fl* animals after 4-months of age and accounted for 10% of total live cells in tumors from 7-month old animals (Figure 3J). Fibroblasts are a major producer of collagen, and collagens account for 80% of the dry weight of neurofibromas (36). To determine if the increase in CD44+;CD49f-fibroblast cell population correlates with increased collagen we analyzed Sirius red staining. Sirius red staining increased in *DhhCre;Nf1fl/fl* mice at 2 months when compared to control, likely reflecting collagen released by *Nf1* mutant Schwann cells and/or resident fibroblasts. A further additional increase in collagen occurred at 4-months, coinciding with tumor onset and formation of CD44+;CD49f-cancer associated fibroblasts (Figure 3, K-L).

Hyaluronic acid (HA), a component of the neurofibroma matrix (56) is a ligand for CD44 (57). HA expression can be determined by staining for HA-binding protein (58). Increased HABP staining is observed after 4-months of age (Figure 3, M-N) concurrent with the elevated CD44+;CD49f-fibroblasts. We also quantified SPP1/osteopontin, another ligand of the CD44 receptor which is transcriptionally elevated in neurofibroma Schwann cells (34,49,59). SPP1 modestly increased in expression at 4-months and further increased at 7 months in *DhhCre;Nf1fl/fl* mice versus control (Figure 3, O-P). Thus, both HA and SPP1 elevation correlate with increase in CD44+;CD49f-fibroblasts in plexiform neurofibroma development.

### NF-κB pathway activation occurs at tumor onset

We posited that CD44+;CD49f+ Schwann cells in tumors are transcriptionally reprogrammed. To define transcriptional alterations in *Nf1-/-*Schwann cells that occur concomitant with CD44+;CD49f+ co-expression we FACS sorted lineage+ cells from wildtype mouse DRG and from *DhhCre;Nf1fl/fl* mice (from tumor and non-tumor DRG) and performed RNA-Seq on the sorted cells. Schwann cell identity was confirmed by expression of *Mbp, Plp1, Pmp22, Gfap, L1cam, Cdh19* and other Schwann cell marker genes (Supplemental Figure 2). Lineage+ tumor Schwann cells (Figure 4A) showed significant increases in “Inflammatory response”, “KRAS signaling”, “Interferon gamma response”, “NF-κB signaling” and “Epithelial to mesenchymal transition/plasticity” associated transcriptional profiles when compared to non-tumor Schwann cells. Transcriptional upregulation of EMP genes (foldchange>2, p-value< 0.05), including numerous collagens (*Col1a1, Col1a2, Col6a2, Col12a1, Col5a2,* and *Col4a2*) was observed in the lineage+ Schwann cells from tumor versus non-tumor DRG or wildtype DRG (Figure 4B). Upregulation of NF-κB target genes (https://esbl.nhlbi.nih.gov/Signaling-Pathways/NF-κB-Targets) was also observed in lineage+ tumor Schwann cells (Figure 4C).

**Figure 4.**
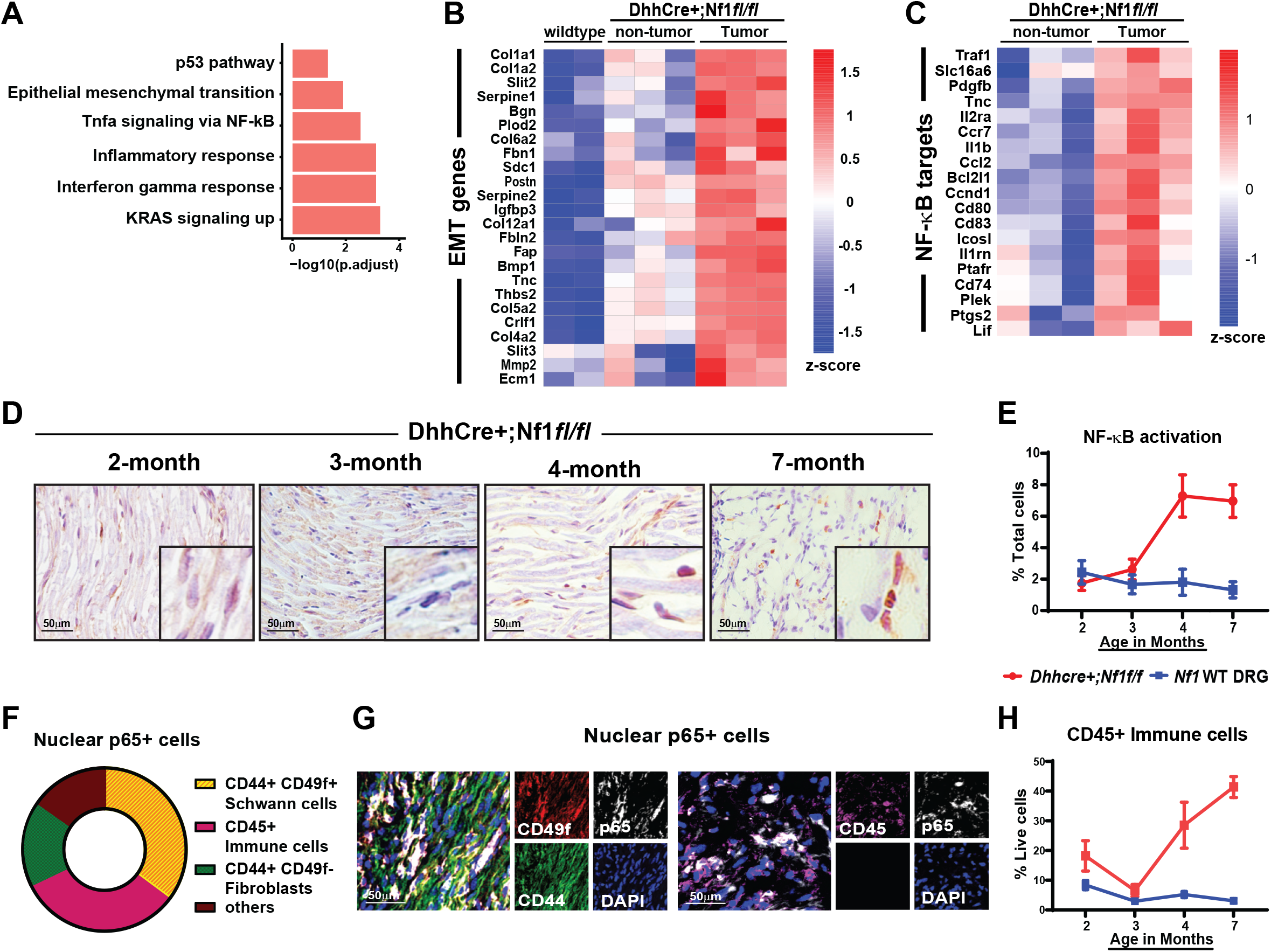
Cellular and extracellular remodeling at tumor onset coincides with enhanced NF-κB pathway activation in plexiform neurofibroma. (A) Hallmark gene sets upregulated in lineage+ Schwann cells flow sorted from *DhhCre+;Nf1fl/fl* tumor compared to non-tumor DRG (foldchage>2, *p-value*<0.05). (B) Differential gene expression analysis of EMT markers in lineage+ tumor Schwann cells compared to Schwann cells isolated from non-tumor DRG and from WT DRG (foldchage>2, *p-value*<0.05). (C) NF-κB target genes expressed in lineage+ tumor Schwann cells compared to non-tumor DRG Schwann cells (foldchage>2, *p-value*<0.05). (D) Representative IHC images for p65 staining of DRG/tumor. NF-κB pathway activation (E) was quantified as %nuclear p65 per HPF. Scale bars, 50μm (n=3 per group). *p*<0.01 for 4- and 7-months. (F) Doughnut plot showing proportions of cells expressing nuclear p65 and (G) representative images of p65 nuclear expression in CD44+ CD49f+ Schwann cells and in CD45+ immune cells. Scale bars, 50μm (n=3-4 per group). (H) Quantification of immune cells in DRG/ tumor by flow cytometry in *DhhCre+;Nf1fl/fl* mice. Total immune cells (CD45+) was quantified as %Live cells (n=3-4 per group). *p*<0.0006 for 4- and 7-months. Significance determined with Ordinary two-way ANOVA and Tukey’s multiple comparisons test. See also Supplemental Figure 2.

This finding was consistent with the hypothesis that NF-κB activation acts as a step in PN formation, contributing to cellular reprogramming and cytokine production. To test this hypothesis, we tested if nuclear p65 expression precedes tumor formation. Rare cells expressing nuclear p65 were present at 3 months of age, and a significant increase in nuclear p65 expressing cells was seen abruptly at the 4-month old time point, when tumors form (Figure 4D-E). Double labeling of tumor tissue sections showed that of the p65+ cells in PN, 35% were CD44+;CD49f+ Schwann cells, 20% were CD44+;CD49f-fibroblasts, and 32% were CD45+ immune cells; (Figure 4, F-G). Consistent with the idea that NF-κB pathway activation-driven cytokine expression recruits immune cells into the tumor microenvironment (38), immune cells (CD45+) also increased at 4-months in the tumor (Figure 4H). These data show that NF-κB signaling in Schwann cells correlates with tumor formation and inflammation.

### Active NF-κB signaling in Schwann cells causes cellular reprogramming

The correlation between tumor formation and nuclear p65 expression in CD44+;CD49f+ Schwann cells suggested that NF-κB activation causes reprogramming of Schwann cells, alters Schwann cell viability, and/or has non-cell autonomous roles. We therefore tested if direct activation of NF-κB signaling in cultured embryonic primary mouse Schwann cells increases CD44+;CD49f+ expression. We infected *Nf1-/-*Schwann cells with active IKK2, which significantly increased CD44+;CD49f+ marker expression (Figure 5A and Supplemental Figure 3, A-B). Fold change in CD44+CD49f+ expression is shown in Supplemental Figure 3A; CD44+;CD49f+ cells represented 0.74% of *Nf1+/+* cells and 38.7% of *Nf1-/-*cells expressing IKK2^CA^ (Supplemental Figure 3A). IL1β, known to activate NF-κB signaling, similarly increased CD44+;CD49f+ co-expression on *Nf1-/-*Schwann cells, with less effect on wild type cells and as compared to other factors (Figure 5B & Supplemental Figure 3C). Cell proliferation remained unaffected by treatment of *Nf1-/-*Schwann cells with IL1β (Supplemental Figure 3D). Serum depletion, which induces cell stress, similarly resulted in increased marker co-expression in *Nf1-/-*Schwann cells with significantly less effect on wild type cells (Figure 5C) and concurrent with significantly increased nuclear phospho-p65 expression (Figure 5D). NF-κB-mediated transcription is predicted to increase transcription of target genes, and subsequent release of cytokines. We tested if increased cytokine release occurs after NF-κB pathway activation in this setting by culturing cells in serum-free medium to activate NF-κB and then subjecting serum-free cell-culture medium to cytokine analyses. Of 22 tested factors (Supplemental Table 2), immune cell chemoattractants (CCL2 and CCL3) increased in the culture medium with significantly higher expression in *Nf1-/-*Schwann cells (Figure 5, E-F). Also, EMP-associated gene expression (*Tnc*, *Snai2*, *Ereg*, and *Serpine1*) increased in stressed serum-starved *Nf1-/-*Schwann cells (Figure 5G), but not in *Nf1+/+* wildtype Schwann cells. Thus, IL1β and serum deprivation are stressors that can activate NF-κB in vitro, correlating with increased cytokine release and increased expression of EMP related genes in primary Schwann cells, effects enhanced by loss of *Nf1*.

**Figure 5.**
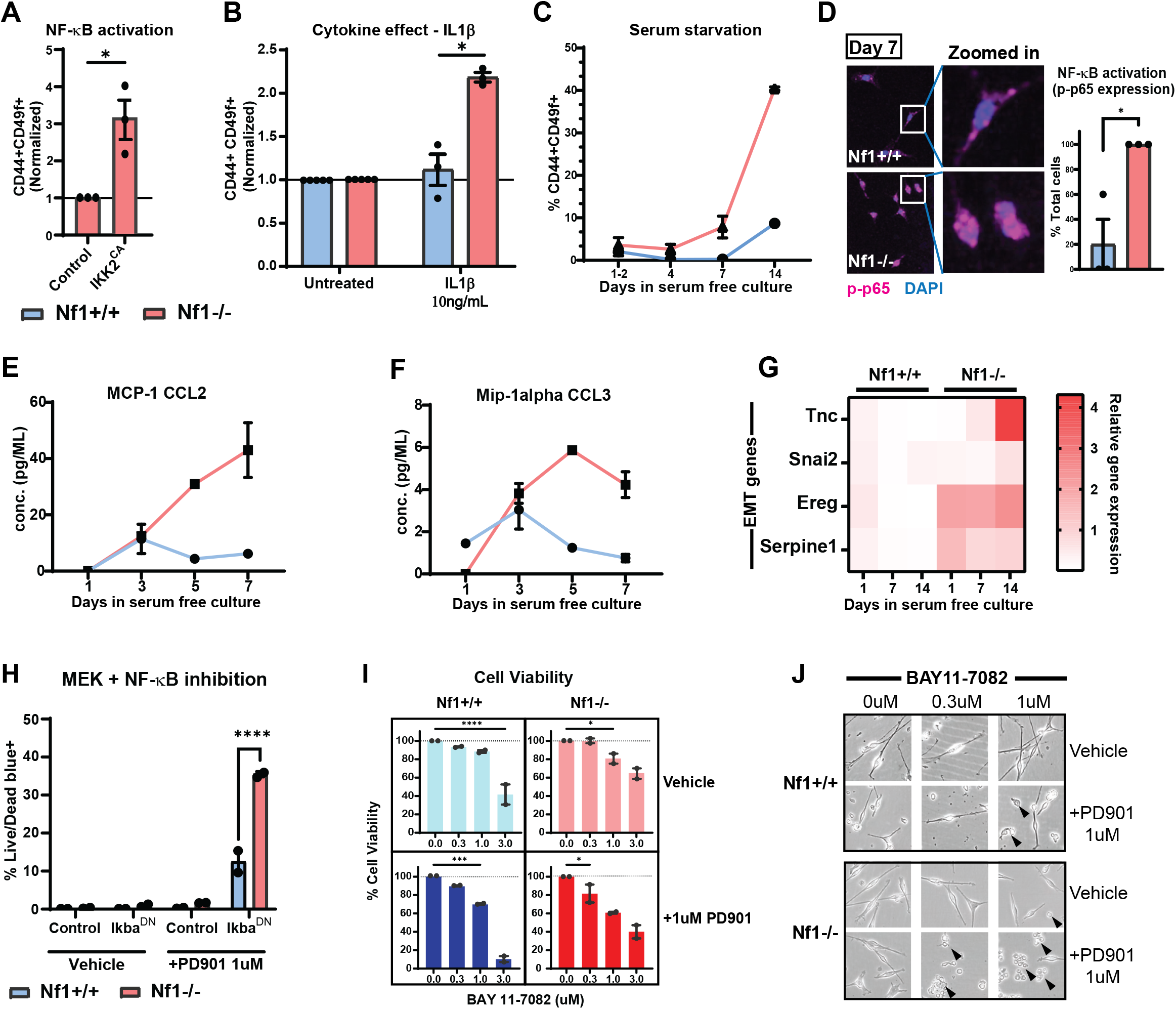
Active NF-κB signaling in Schwann cells causes cellular reprogramming *in-vitro*. (A) Bar plot of constitutively active NF-κB signaling (IKK2^CA^) on CD44+ CD49f+ coexpression on Nf1-/- primary mouse Schwann cells. *p*<0.01, Unpaired t-test. (B) Quantification of CD44+ CD49f+ coexpression on primary mouse Schwann cells after 24-hour treatment with 10ng/mL IL1β. *p*<0.01, Unpaired t-test. (C) Change in CD44+ CD49f+ expression on Schwann cells quantified after prolonged serum starvation (stress) through days 1-14. *p*<0.005 for 7- and 14-day comparison, Ordinary two-way ANOVA. (D) Representative image of phospho-p65 staining in primary mouse Schwann cells at day 7 in serum starved conditions. Quantification on right. *p*<0.0001, Unpaired t-test. (E) Change in expression of chemoattractants CCL2 and CCL3 (F) produced by stressed Schwann cells and quantified by analysis of conditioned media. *p*<0.01 for 7- and 14-day comparison, Ordinary two-way ANOVA. (G) Relative mRNA quantification (to Nf1+/+ Day1) of EMT markers expressed by primary mouse Schwann cells under stress. (H) Bar plot of cell death (%Live/Dead Blue+) in primary mouse Schwann cells expressing Ikba^DN^ (supre-repressor of NF-κB signaling) and treated with MEK inhibitor PD0325901 (PD901). *p*<0.0001, Unpaired t-test. (I) Quantification of cell viability upon treatment of serum starved primary mouse Schwann cells with NF-κB pathway inhibitor BAY11-7082 as a single agent and in combination with PD901. (J) Representative images of Nf1-/- and Nf1+/+ Schwann cells after treatment with BAY11-7082 and PD901. Arrows mark the dead cells. See also Supplemental Figure 3 and Supplemental Table 2.

NF-κB signals either through a canonical pathway, resulting in translocation of p65/RelA to the nucleus, or a non-canonical pathway that drives p52/RelB to the nucleus. To test if either or both pathways have a cell autonomous role in Schwann cells, we blocked NF-κB signaling using inhibitors of canonical or noncanonical NF-κB signaling (60) and monitored cell viability and expression of CD44+;CD49f+ by flow cytometry. Three inhibitors that block the canonical NF-κB pathway (BAY11-7082, UC65, and parthenolide) each inhibited cell-viability in a dose-dependent fashion and reduced CD44+;CD49f+ co-expression in *Nf1-/-*Schwann cells (Supplemental Figure 3, E-H). These compounds target Ikba directly or Ikba ubiquitination, thereby preventing p65 movement to the nucleus and canonical NF-κB pathway activation. These inhibitors can also affect additional cell-signaling pathways; BAY11-7082 can act as an inhibitor for protein tyrosine phosphatase (PTP) (61) and BAY11-7082 and parthenolide block the NLRP3 inflammasome pathway (62). Upstream inhibitors of the canonical pathway (inhibitors of TAK1, or IRAK4) had no effect as single agents (Supplemental Figure 3, E-G), suggesting that multiple and/or as-yet unidentified inputs activate NF-κB in Schwann cells. TBK1 is recruited as a response to cGAS/STING pathway activation (Supplemental Figure 3E), which is a contributor to neurofibroma formation (35); however, blocking TBK1 activity did not affect Schwann cell viability (Supplemental Figure 3, F-G). A NIK inhibitor, which inhibits the non-canonical NF-κB pathway, also had no effect on Schwann cell viability (Supplemental Figure 3, E-G). We tested the effect of BAY11-7082, UC65 and parthenolide on *Nf1+/+* Schwann cells under the same conditions and show that these inhibitors do not affect cell viability or CD44+;CD49f+ expression in *Nf1*-wildtype Schwann cells (Supplemental Figure 3H(b)). These data support the hypothesis that NF-κB activation can control viability of *Nf1-/-*Schwann cells, and that expression of CD44 in CD49f expressing Schwann cells can be driven by canonical NF-κB pathway activation.

We next tested whether combining MEK inhibition with NF-κB blockade alters *Nf1-/-*Schwann cell growth in vitro. First, we blocked NF-κB signaling in primary mouse Schwann cells with a Ikba^DN^ super-repressor, which shows more specificity than available inhibitors. We tested Ikba^DN^ super-repressor alone or in combination with 1uM MEK inhibitor PD0325901/Mirdametinib. *Nf1-/-*Schwann cells showed increased cell-death on live/dead blue staining with the super repressor infection, while *Nf1+/+* Schwann cells showed less effect (Figure 5H). We then tested if combining MEK inhibition (1uM) with NF-κB inhibitor BAY11-7082 would also cause cell death. This combination decreased viability by 20% (Figure 5I) at a lower concentration of BAY11-7082 (0.3uM) in *Nf1-/-* Schwann cells, and an increase in cells with the morphology of dying cells was observed on phase microscopy (Figure 5J). This effect was slightly enhanced in *Nf1-/-* versus wild type Schwann cells. This result suggested that combination therapy might affect established tumors.

### Dual inhibition of NF-κB and MEK signaling pathways targets Schwann cell survival in established plexiform neurofibroma

BAY11-7082 is a NF-κB inhibitor that inhibits Ikba ubiquitination and thus prevents p65 nuclear translocation (63–66). We treated tumor bearing *DhhCre;Nf1fl/fl* mice for 2-weeks with various doses of BAY11-7082, with an endpoint dose administered 2 hours before euthanasia, then analyzed nuclear-p65 expression and cleaved caspase-3 expression. Staining of tissue sections confirmed significantly reduced nuclear p65 expression at the 10mg/kg dose of BAY11-7082 (Figure 6A). Cell-death was elevated at all doses tested as indicated by CC3 staining (Supplemental Figure 4A). Tumor-bearing mice were subjected to an MRI scan at 5 months to measure tumor size and at 7 months to establish tumor growth rate. Treatment started at 7 months and continued for 2 months. Mouse weight was monitored biweekly. At the 9-month endpoint, combination therapy, like single agent MEK inhibitor shrunk the tumors (Figure 6B). The extent of tumor shrinkage of the combination was not significantly greater (*p* value = 0.6017) than with MEK inhibitor alone and was not durable. Despite the failure of this agent to shrink tumors, there were changes within established tumors treated with the combination. Staining of tissue sections revealed a significant reduction in cell proliferation (Figure 6C) and NF-κB signaling (nuclear p65) (Figure 6D) in the combination treatment group. While only combination treatment significantly reduced cell proliferation (Figure 6C), both single agents as well as the combination treatment had significant effects on NF-κB signal inhibition, with combination having <1% cells with nuclear p65 expression at treatment endpoint. On flow cytometric analysis, in the NF-κB and MEK inhibitor combination treatment cohort, there was a significant reduction in the proportion of CD45+ immune cells, CD44+;CD49f+ Schwann cells, and CD44+;CD49f-fibroblasts (Figure 6, E-F). In contrast, as a single agent, MEK inhibitor treatment significantly reduced the CD44+;CD49f+ Schwann cells but not the immune cells or the fibroblasts. The NF-κB inhibitor as a single agent significantly reduced the CD44+;CD49f-fibroblast population but not the Schwann cells or the immune cells (Figure 6, E-F). Lineage+ *Nf1* mutant Schwann cells were also significantly reduced after two months of combination treatment, but not with single agent treatments (Figure 6G). Cytokine profiling of tumor tissue lysates confirmed that NF-κB inhibition both as a single agent and in combination treatment counteracted pro-inflammatory mediators (C5a, ICAM1) elevated by MEK inhibition alone (Figure 6H and Supplemental Figure 4B). Although the combination of NF-κB inhibitor treatment with the MEK inhibitor did not shrink established tumors better than single agent MEK treatment, the number of tumors present upon gross dissection at the 2-month treatment endpoint was significantly reduced in the combination treatment cohort when compared to both single agents (Figure 6I). Notably, tumors that formed in cervical regions were absent, with fewer tumors present in the thoracic regions (Figure 6J). These results were consistent with the hypothesis that NF-κB signaling plays a role in plexiform neurofibroma initiation. In contrast, while NF-κB controls significant aspects of tumor biology in established tumors, pathway blockade does not enhance the effect of MEK inhibition on tumor shrinkage in established tumors.

**Figure 6.**
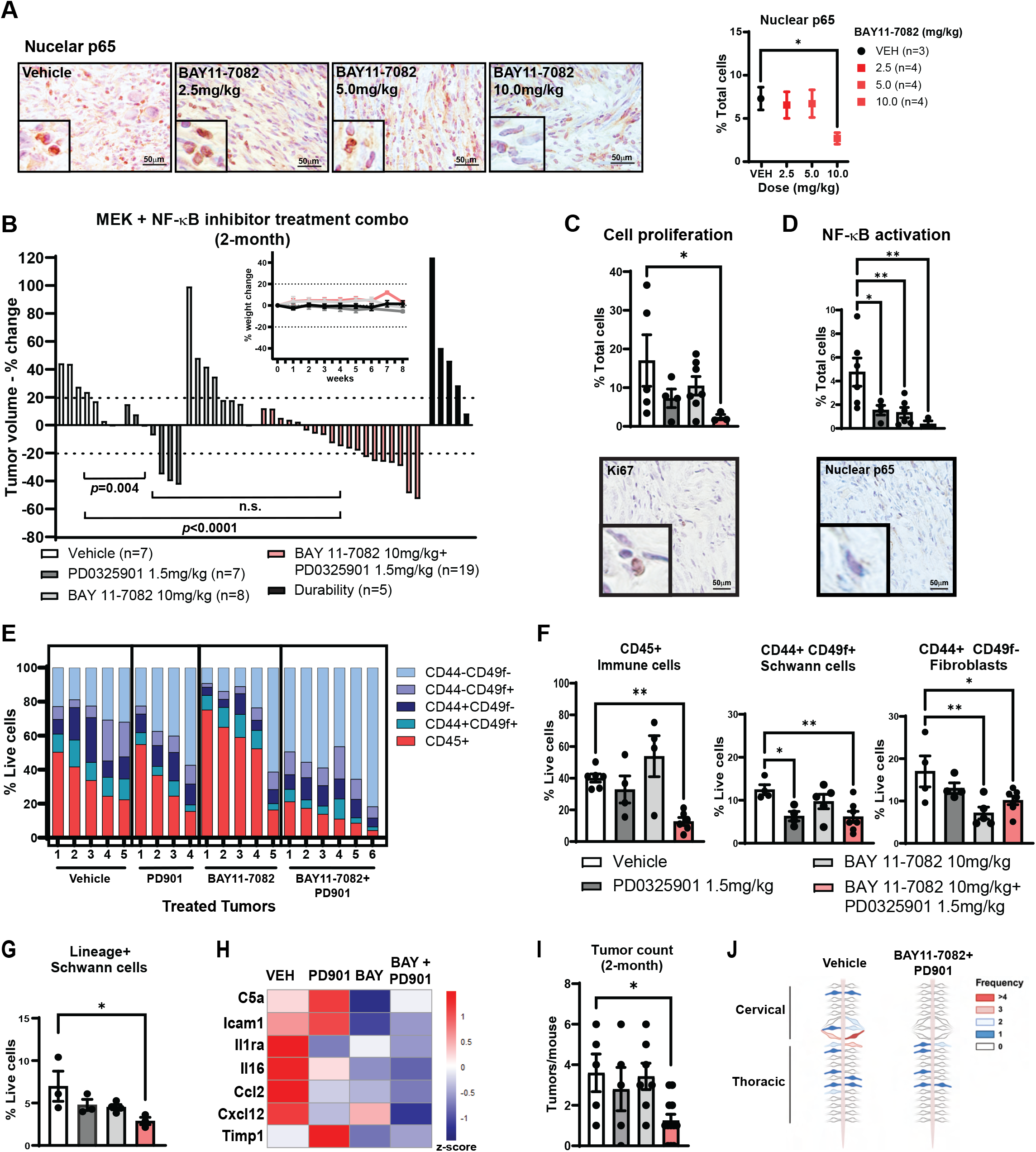
Dual inhibition of NF-κB and MEK signaling pathways targets Schwann cell survival. (A) Representative images of nuclear p65 expression on 2-week treatment with BAY11-7082 (left) and quantification of percent expression (right). (B) Waterfall plot of tumor volume change in *DhhCre+;Nf1fl/fl* mice after 2-month combination treatment with BAY11-7082 and PD901. Each bar represents percent volume change for a single mouse (Significance determined by random effects model analysis). Inset: Mouse weights plotted over 2-month treatment period. Weights plotted relative to day 0. (C) Immunohistochemical analysis of cell proliferation quantified as percent Ki67+ cells per tumor (top) after 2-month combination treatment; representative image of Ki67 staining in the BAY11-7082 + PD901 combination treated mice (bottom). Scale bars, 50μm. (D) Quantification of nuclear p65 expression after 2-month treatment (top) and representative image of nuclear p65 staining of combination treated tumors (bottom). Scale bars, 50μm. (E) Stacked bar plots per sample clustered by percentage of each identified subpopulation by flow cytometry. (F) Change in CD45+ Immune cells (left), CD44+;CD49f+ Schwann cells (middle) and CD44+ CD49f-Fibroblasts (right) quantified by flow cytometry analysis of tumor tissue at endpoint of 2-month treatment conditions. (G) Lineage+ Schwann cells quantified after 2-month treatment by flow cytometry. (H) Heatmap of row normalized expression of assayed cytokines and proteins in treated tumor tissue. (I) Quantification of tumor number count after 2-month treatment. (J) Map of tumor-specific location on DRG/Nerve regions after 2-month treatment Significance unless specified is determined by Ordinary one-way ANOVA. \**p*<0.0332, \*\**p*<0.0021, \*\*\**p*<0.0002, \*\*\*\**p*<0.0001, ns [non-significant]. See also Supplemental Figure 4.

### NF-κB blockade prevents PN formation in vivo

Based on these findings, and because NF-κB activation occurs concurrent with tumor formation, we hypothesized that blocking NF-κB activation pre-tumor formation (before 4-months) would delay or prevent tumor growth. To test this, 3-month-old *DhhCre;Nf1fl/fl* mice were treated with 10mg/kg of BAY11-7082 for 1-month. Then, the mice were left untreated for 2.5 months (Figure 7A). Gross dissections of spinal cords and attached nerves are shown in Figure 7B. We counted tumor number and measured tumor size in these preparations. Tumor number/mouse at 6.5 months of age in vehicle (experimental control) treated mice was consistent with historical tumor counts at this age in the *DhhCre;Nf1fl/fl* mice (22). Strikingly, 3/7 BAY11-7082 treated mice showed no tumors, and an average of n=1 tumor/mouse, a significant reduction in tumor number compared to both experimental (69%) and historical controls (75%) at the same time point (Figure 7C (left)). DRG/tumor diameter in treated mice was also significantly reduced (Figure 7C (right)). Staining of tissue sections from these mice verified significantly reduced NF-κB (Figure 7D) and CD44+;CD49f+ Schwann cells in DRG/nerve lacking tumor (Figure 7E) at the end of the experiment, consistent with the near-absence of tumor formation. Flow cytometry analysis verified a reduction in CD44+;CD49f+ Schwann cells both as a percent of the total live cells and as a percent of the total Dhh-lineage+ cells in these treated non-tumor DRG/nerve samples (Figure 7, F-G). Of the remaining CD44+;CD49f+ Schwann cells, a downward trend in the expression of mesenchymal markers was observed, with CD34+ expression showing a significant reduction compared to control (Figure 7H (left)). The CD45+ immune cells were also reduced in the treatment group (Figure 7H (right)). Thus, transient blockade of NF-κB signaling prior to tumor onset significantly restricts PN formation.

**Figure 7.**
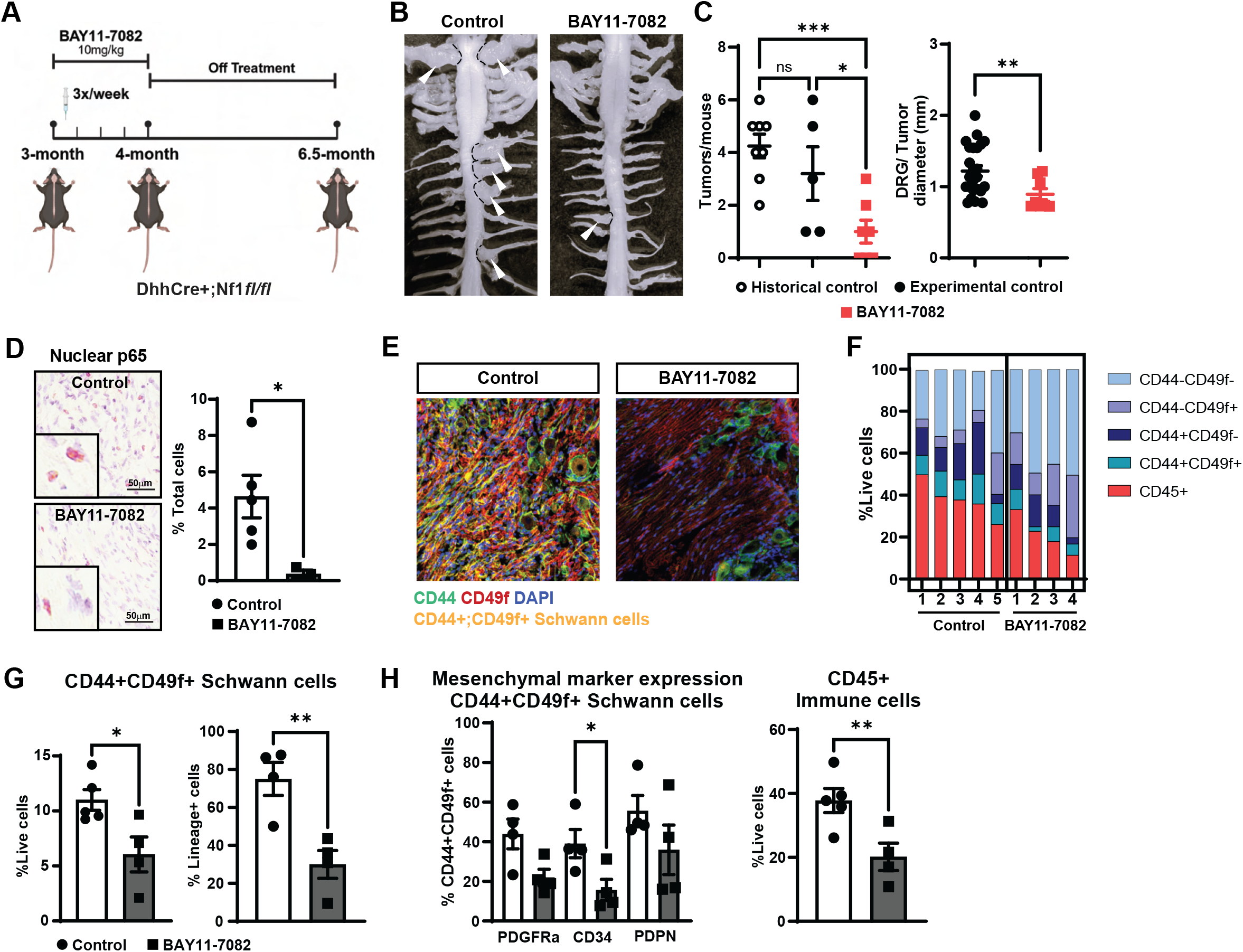
Blocking NF-KB activation delays neurofibroma onset in mice. (A) Schematic overview of NF-KB pathway inhibition pre-tumor formation (3-4months of age). (Created with BioRender.com) (B) Representative image of gross dissection of spinal cord and tumor/DRG comparing vehicle treated and BAY11-7082 inhibitor treated mice at 6.5 months of age. (C) Tumor number quantification per mouse (\**p-value* = 0.0259, \*\*\**p-value* = 0.0008, ns [non-significant] determined by Ordinary One-Way ANOVA) and DRG/ tumor diameter (in mm) quantification (\*\**p*<0.01, Welch’s t-test) at 6.5 months of age after 4-week NF-KB inhibitor treatment (BAY11-7082, 10mg/kg) at pre-tumor stage (3 months). (D) Image of nuclear p65 expression in BAY11-7082 inhibitor treated tissue vs control at 6.5 months. Scale bars, 50μm. Quantification of nuclear-p65 expression on right. \**p*<0.05, Welch’s t-test. (E) Representative images of CD44+;CD49f+ Schwann cells at 6.5 months of age after NF-KB inhibitor prevention treatment. (F) Stacked bar plots of flow cytometry analyzed data clustered by percentage of identified subpopulations in BAY11-7082 inhibitor treated group at 6.5 months of age and the control group. (G) Change in Schwann cell population quantified by flow cytometry analysis as a percent of live cells (left) and as a percent of Dhh-lineage+ cells (right) at 6.5 months of age. \**p*<0.05, \*\**p*<0.01, Welch’s t-test. (H) Quantification of mesenchymal marker expression by flow cytometry analysis on CD44+;CD49f+ Schwann cells, and CD45+ immune cells after NF-KB inhibitor prevention treatment. (\**p*<0.05, \*\**p*<0.01, Welch’s t-test)

## Discussion

The absence of cooperating genomic lesions in plexiform neurofibroma Schwann cells has long implied that non-mutational events can drive tumorigenesis. The identity of such events has remained unknown. This study aimed to identify the mechanism(s) that regulate tumor onset to inform the development of therapeutics targeted at preventing tumor formation in individuals with Neurofibromatosis Type I disease. Our findings identify NF-κB activation as a critical, targetable step in plexiform neurofibroma initiation. We identified CD44 and CD49f, and additional markers of Schwann cell reprogramming/EMP on tumor Schwann cells and found that NF-κB activation in Schwann cells increases marker expression. These markers are absent on normal Schwann cells. We show that NF-κB activation is critical for tumor initiation after *Nf1* loss, as transient blockade of NF-κB activation prior to tumor formation significantly restrains tumor initiation.

We identified CD49f and CD44 on the surface of tumor Schwann cells by spatial analysis and verified expression by flow cytometry. Double positive cells were abundant in tumors but not pre-tumors. CD49f/integrin alpha 6 is encoded by the *Itga6* gene and CD49f is a known Schwann cell surface protein. In Schwann cells, ITGA6/CD49f associates with integrin beta 1 or integrin beta 4 subunit to form complexes that interact with extracellular matrix proteins, including the laminins which are abundant in neurofibromas (37,68). Of note, in the Schwann cell basal lamina, laminins inhibit PKA-mediated Nrg1-III signaling via ErbB2/ErbB3 to prevent myelination until radial sorting is complete (69), activate the co-transcriptional activators YAP1 and TAZ to induce cell proliferation, increase dystroglycan and ITGA6B1 expression, and increase myelination-driving Egr2/Krox20 transcription factors (70). We note that expression of CD49f, as read out by flow cytometry, does not change on Schwann cells in the absence of NF1. In contrast, CD44 increases in tumor Schwann cells. CD44 is a cell-surface hyaluronic acid (HA) receptor, a glycoprotein involved in cell-cell interactions, cell adhesion and migration (49). CD44 interacts with extracellular ligands including osteopontin/SPP1, collagens, and matrix metalloproteinases (MMPs) (49,67), the expression of which increase in plexiform neurofibroma Schwann cells. HA and osteopontin are present in the plexiform neurofibroma environment, and both are elevated after tumor onset when fibroblasts in the tumor increase. Therefore, while we cannot exclude HA and osteopontin as ligands for CD44 increase in plexiform neurofibroma, its relevance and potential to elevate expression of CD44 during tumor onset need to be studied. Whether CD44+ simply marks tumor Schwann cells or whether signaling through CD44 has functional importance to tumor formation warrants further investigation in this setting.

A recent study highlighted a glial-mesenchymal transcriptional transition that occurs during the progression of neurofibroma to aggressive MPNST (71). Our analysis indicates that the increased expression of mesenchymal genes and proteins occurs in Schwann cells earlier, as PNs form. Transcriptional profiling of flow sorted lineage+ tumor Schwann cells showed significant enrichment of a mesenchymal gene signature, and further analysis confirmed the expression of the mesenchymal markers PDGFRα and podoplanin on lineage+ tumor Schwann cells. In human PNs, the expression of the mesenchymal marker CD34 on p75+;CD44+;CD49f+ Schwann cells underscores the relevance of this finding across species. Of note, since the 1940s it has been debated whether neurofibroma tumor cells are fibroblasts or Schwann cells (72). The findings that human neurofibroma cells show a continuous Schwann cell basal lamina (73) and that neurofibroma Schwann cells but not fibroblasts stimulate angiogenic reactions (74), together with the finding that loss of *Nf1* in mouse Schwann cells is sufficient to drive neurofibroma formation, all support the currently accepted idea that Schwann cells are the neurofibroma cells-of-origin. Together with our findings, *NF1* mutant Schwann cells acquire mesenchymal characteristics, helping to reconcile the early findings that neurofibroma cells can show intermediate morphological phenotypes (75). Our results are consistent with acquisition of mesenchymal traits in Schwann cells after nerve injury (76) and with the demonstration that Glut1, normally a nerve perineurial cell marker, becomes detectable in injury-induced neurofibroma Schwann cells (77).

This study demonstrates that inflammatory cues including IL1β and cellular stress activate NF-κB signaling in cultured Schwann cells, reprogramming them into the CD44+;CD49f+ tumor Schwann cells present within plexiform neurofibroma. Our model aligns with previously published literature linking NF-κB as a regulator of chronic inflammation and tumor development (38,78,79). We have not yet identified the in vivo signals that activate NF-κB signaling in *Nf1-/-*Schwann cells. These are likely to include oncogenic cellular stress that results in immune cell infiltration and matrix alterations, caused by the presence of *Nf1-/-* Schwann cells. NF-κB-activated genes such as *Cxcl10* are expressed early in plexiform neurofibroma formation (29). EGFR, shown to disrupt nerve structure (80) and increase neurofibroma progression (19) can activate NF-κB (81,82). IL1β is another candidate factor; IL1β exposure increased the CD44+;CD49f+ expression on Schwann cells and is produced by neurofibroma macrophages; IL1 receptors are present on Schwann cells (34,44). An IL1β-IL1R feed forward loop would have the potential to amplify NF-κB signaling.

Whichever ligand(s) drive signal activation, the result of NF-κB activation is likely to be immune cell recruitment and reprogramming of fibroblasts in the environment given that the release of the immune cell chemoattractants CCL2 and CCL3 is induced by NF-κB activation in vitro, and that expression of immune modulators decreased after blocking NF-κB signaling in established tumors. In normal Schwann cells, blocking NF-κB activation in GFAP expressing non-myelinating Schwann cells reduced inflammatory immune cell chemoattractants including CCL2 and CXCL10 after sciatic nerve injury (41,42). Importantly, while it was initially suggested that interfering with NF-κB signaling in Schwann cells affects normal cell differentiation, later reports (83) demonstrated that NF-κB is not required and that blocking NF-κB only modestly delays timely remyelination following peripheral nerve injury (84). We are therefore encouraged that transiently or intermittently blocking the NF-κB pathway might be feasible as a neurofibroma prevention strategy.

NF1 loss in cells results in elevated intracellular RAS-GTP; later, NF-κB activation is necessary for tumor onset, as evidenced by the substantial reduction in tumor formation when predisposed mice were treated with NF-κB inhibitor for one month prior to tumor formation. We conclude that elevated MAPK signaling in *Nf1-/-* Schwann cells is necessary for sustained CD44+;CD49f+ co-expression, based on our finding that treating established tumors with MEK inhibitors, but not NF-κB inhibitors, reduced the CD44+;CD49f+ tumor Schwann cell population. Also, in vitro, blocking MEK reduced CD44+;CD49f+ Schwann cells. CD44+;CD49f+ expression was increased when NF-κB signaling became active in Schwann cells, especially in *Nf1-/-* Schwann cells. Collectively, this data suggests that active MEK signaling lowers the threshold necessary for NF-κB signaling to increase the CD44+;CD49f+ co-expression on the Schwann cells.

Based on our tumor prevention data and the reduction of tumor Schwann cells, immune cells and fibroblasts with combination treatment, we expected to see a durable tumor volume reduction in MEK and NF-κB combination treated mice. The lack thereof suggests that additional, unidentified, pathways are in play once tumors are established. A candidate pathway is STAT3; STAT3 inhibition in Schwann cells reduces tumor numbers in the DhhCre mouse model (85) and its activation in PNs increases the macrophage cell populations (43) However, it remains possible that NF-κB and STAT3 have non-redundant roles to play in the PN inflammatory environment. It will be important to better understand the interactions among signaling pathways to extensively and durably shrink neurofibromas.

The natural history of the growth of PN in human NF1 is well described. Young children (age ≤8 years) have the fastest PN growth rate (86) and there is no acceleration in growth rate during puberty (87). In adults, there is minimal PN growth and even small decreases in PN volumes over time (<20% tumor shrinkage) (88,89). These findings have suggested that an ideal treatment strategy may be to prevent tumor growth and the development of morbidity. This hypothesis is currently being tested using the MEKi selumetinib (NCT06188741). Also, in humans cutaneous neurofibromas develop later, largely at puberty. These tumors, which show little morbidity but can be highly disfiguring, might also benefit from early intervention. It will be of significant interest to determine whether NF-κB is activated in these cutaneous tumors as it is in plexiform neurofibromas. Although no NF-κB inhibitors are currently FDA-approved, agents targeting this pathway, including derivatives of parthenolide such as ACT001 are under clinical evaluation in certain malignancies (91). In our study in cultured Schwann cells, parthenolide showed similar effects to BAY11-7082. Another recent study identified DMAPT, an orally bioavailable parthenolide analog, as a compound that reduced growth of immortalized NF1-/- Schwann cells (92). The availability of NF-κB inhibitors in the clinic is hoped to pave the way for exploratory prevention trials in children predisposed to Neurofibromatosis Type 1 disease.

## Materials and Methods

### Mice

DhhCre+ Nf1fl/fl mice were bred in-house at Cincinnati Children’s Hospital Medical Center (93). All procedures were approved by the Institutional Animal Care and Use Committee. Mice were housed under controlled temperature and humidity, with free access to food and water, on a 12-hour light/dark cycle. Both male and female mice were used.

### Multiplexed Spatial Imaging

#### Tissue preparation

Fresh frozen plexiform neurofibroma tissue samples were collected at 7 months of age and embedded in OCT compound (FisherScientific, Cat. No. 23–730–571). Tissue sections (10 µm thickness) were cut onto poly-L-lysine coated coverslips (Electron Microscopy Sciences, Cat. No. 72204-01). Coverslips were manually coated with 0.1% poly-L-lysine solution (Santa Cruz Biotechnology, Cat. No. sc-286689A) prior to use.

#### Antibody panel and staining

A 27-plex mouse antibody panel was optimized for spatial analysis of the plexiform neurofibroma environment. Tissue staining and imaging were performed using the Phenocycler (formerly CODEX) Ultra-High Parameter Imaging system at the NCI Imaging Core, Bethesda, Maryland (94). Sections were stained with iterative annealing and stripping of fluorescently labeled oligonucleotide-barcoded antibodies. Cyclic image acquisition was performed using four filters (DAPI, FITC, CY3, CY5) on a Keyence BZ-X810 microscope.

#### Image processing and analysis

Images were processed for pixel-precise alignment, background subtraction, stitching, and quality improvement, generating TIFF files for downstream analysis. Halo digital pathology software (Highplex FL module) was used for segmentation and single-cell spatial analysis. Parameters included Hoechst+ nuclei segmentation, maximum cytoplasm radius (1.5 µm around each nuclei), membrane segmentation aggressiveness (0.8), and cell size (min = 5 µm², max = 300 µm²). Marker intensity thresholds were visually defined to determine the minimum cutoff for each object to be considered positive for that marker, with blanks included to exclude non-specific staining artifacts. High-resolution multiplexed CODEX fluorescence images can be interactively explored via the HALO Link Viewer (https://halo.cancer.gov/s/studies/19261). The full raw image TIFF files along with mapped single-cell segmentation masks and raw marker signal intensity values, are available from the corresponding author upon reasonable request.

#### RNAseq analysis

The libraries were prepared with polyA selection using the Truseq RNA Library Prep Kit, and the libraries were sequenced at an average depth of 50M paired-end 150bp nucleotide reads. After the quality of reads was examined using FastQC (v0.11.7, https://www.bioinformatics.babraham.ac.uk/projects/fastqc), paired-end reads were aligned against mouse mm10 genome (https://support.illumina.com/sequencing/sequencing_software/igenome.html) using HISAT2 (v2.0.5, http://daehwankimlab.github.io/hisat2). The raw gene counts were calculated using featureCounts (v1.5.2, http://subread.sourceforge.net) and normalized using edgeR (v3.16.5, https://bioconductor.org/packages/release/bioc/html/edgeR.html). Differentially expressed genes were predicted using limma/voom (v3.30.6, https://bioconductor.org/packages/release/bioc/html/limma.html). Raw sequencing data is available through Gene Expression Omnibus (GEO) (GSE346553).

#### Human tumor sample processing

Human PN samples were collected at the Mayo Clinic after medically mandated surgeries. Mayo Clinic’s IRB approved a Waiver for HIPAA Authorization and Informed Consent. Samples were enzymatically dissociated as previously described (19).

#### Flow cytometry

To generate a single cell suspension, the tissues were cut into small (∼1mm^3^) pieces and digested in an enzyme mix containing final concentrations of 10 mg/mL Collagenase A (Sigma-Aldrich, Cat. No. 10103586001), 10mg/mL Collagenase Type 4 (Worthington Biochemical, Cat. No. LS004188), 100ug/mL Soybean trypsin inhibitor (Sigma-Aldrich, Cat. No. 10109886001), 250 units of DNAse I (Sigma-Aldrich, Cat. No. 04536282001) and 5mM CaCl2, in complete RPMI media (ThermoFisher Scientific, Cat. No. 11875-119) at 37°C in a shaking incubator for 30 minutes. After digestion, 10% FBS in RPMI was used to stop the reaction, and the cells were filtered, washed, and pelleted. Cells were then resuspended in flow buffer and counted. 1×10^6^ cells were used for each flow experiment.

Cells were stained for 30 minutes with Live/Dead Blue (FisherScientific, Cat. No. L23105), washed and stained with a panel of antibodies that identify the desired cell populations. Cells were then fixed in 4% paraformaldehyde and analyzed within 24 hours of staining. The Cytek Aurora spectral flow cytometer was used for analysis of stained samples. Downstream gating and analysis was done using the FlowJo software.

The following antibodies were used for flow analysis - CD44 (BD Biosciences, Cat. No. 612799 & BioLegend, Cat. No. 103011), CD49f (BD Biosciences, Cat. No. 563707), CD45 (BioLegend, Cat. No. 103147 & BD Biosciences, Cat. No. 564358), GFAP (BD Biosciences, Cat. No. 561483), p75/NGFR (Alomone Labs, Cat. No. ANT-007-AO & BD Biosciences, Cat. No. 560927), CD34 (BD Biosciences, Cat. No. 562608 & 569043), PDGFRα (BD Biosciences, Cat. No. 568239), Podoplanin (BioLegend, Cat. No. 127407).

#### Cell culture

E12.5 DRGs were isolated from mouse embryos, and primary Schwann cells were cultured as previously described (95). Cells were maintained in DMEM (ThermoFisher Scientific, Cat. No. 11965118) + 10% Fetal bovine serum (+ 10ng/mL bHRG (Peprotech, Cat. No. 100-03-500ug) and 2uM Forskolin (Cayman Chemical, Cat. No. 11018) + 1% Pen/strep (FisherScientific, Cat. No. 15-140-122). Cells were incubated at 37°C and 7.5% CO2 for 5 days, after which cells were plated in 24 well plates at passage 2 for experiments. For serum free culture conditions, N2 media (DMEM+F12 (ThermoFisher, Cat. No. 10565042) + 1x N2 supplement (ThermoFisher Scientific, Cat. No. 17502048) + 10ng/mL bHRG + 2uM Forskolin + 1% Pen/strep) was used.

To obtain Schwann cells expressing Ikba^DN^ and IKK2^CA^, cells were transduced with lentiviral particles. Lentivirus plasmids were under control of the CMV promoter, contained a GFP reporter, and (GSG)P2A linker. To create a control plasmid, we took the backbone from an existing plasmid (Addgene #135459) and inserted a gblock containing a GFP(GSG)P2A-STOP sequence through Gibson assembly. We also developed lentiviruses expressing Ikba^DN^ and IKK2^CA^ by opening our backbone plasmid through restriction enzyme digestion and inserted either Ikba^DN^ “super-repressor” sequence (obtained from Addgene #15291) or IKK2^CA^ sequence (obtained from Addgene #11105) through Gibson assembly. Schwann cells were transduced at an MOI of (add number here), 24 hours after plating. FlowJo analysis for CD44;CD49f marker expression was performed on GFP+ cells.

### Immunostaining

Tissues were harvested from PBS-perfused mice, fixed in 4% paraformaldehyde (Electron Microscopy Sciences, Cat. No. 50-980-488), 30% sucrose and embedded in OCT. Sections were cut to 10 uM thickness. Prior to staining, the sections were air dried, fixed in 4% PFA and then incubated in 3% H₂O₂ (FisherScientific, Cat. No. H325-100) solution to quench endogenous peroxidases. Blocking was performed with 10% bovine serum albumin (GeminiBio, Cat. No. 700-100P) + 10% normal goat serum (Jackson Immuno research, Cat. No. 005-000-121) in 1x PBS for 1-hour followed by primary antibody incubation (10% BSA + 1% normal horse serum + 1x PBS). For immunofluorescence staining, conjugated surface antibodies were incubated for 1-hour; unconjugated nuclear antibodies overnight. Nuclei were stained with DAPI (0.1ug/mL, 5 min) (ThermoFisher Scientific, Cat. No. 62248). Imaging was performed on a Nikon Ti2 Eclipse confocal microscope for immunofluorescence.

The following antibodies were used for immunostaining – HABP (Millipore Sigma, Cat. No. 385911), SPP1 (Cell Signaling Technology, Cat. No. 88742), p65 (Cell Signaling Technology, Cat. No. 8242), phospho-p65 (Cell Signaling Technology, Cat. No. 3033), Ki67 (Cell Signaling Technology, Cat. No. 12202), CC3 (Cell Signaling Technology, Cat. No. 9661), Sirius red (AmsBio, Cat. No. 9046).

### qRT-PCR analysis and primers

Total RNA from in vitro cultures was extracted using the miRNeasy Micro Kit (Qiagen, Cat. No. 217084) according to the manufacturer’s instructions. cDNA was synthesized from using the iScript Advanced cDNA Synthesis Kit (Bio-Rad, Cat. No. 1725037). qRT-PCR was performed using SYBR® Green Supermix (Bio-Rad, Cat. No. 1708882) on a CFX Opus 384 well Real-Time PCR System (Bio-Rad, Cat. No. 12011452). Reactions were performed in triplicate using cDNA as template. Data were analyzed using CFX Maestro 2.3 software, and relative mRNA expression levels were calculated using the ΔΔCt method.

The primer sequences used for mouse genes were as follows:

Tnc (F) 5′-CACCACAGCACTGATCACCT-3′, (R) 5′-TCAGGTCTCAGGTTCCCGAT-3′;

Snai2 (F) 5′-AAGCCCAACTACAGCGAACT-3′, (R) 5′-TGAAGTGTCAGAGGAAGGCG-3′;

Ereg (F) 5′-CGCTGCTTTGTCTAGGTTCC-3′, (R) 5′-CAGTAGCCGTCCATGTCAGA-3′;

Serpine1 (F) 5′-GCTGGTGAATGCCCTCTACT-3′, (R) 5′-TGCCTTTCCACCCAGTCATT-3′;

Actb (F) 5′-TCCAGCCTTCCTTCTTGGGTATG-3′, (R) 5′-CGCAGCTCAGTAACAGTCCG-3′.

### In-vivo treatment studies

Mice were MRI scanned at 5 and 7 months to establish tumor growth prior to treatment. BAY11-7082 (Selleck Chemicals, Cat. No. S2913) (10 mg/kg in 40% PEG300 (Millipore Sigma, Cat. No. 8074841000), 0.05% DMSO, 5% Tween80 in 1x PBS) was administered intraperitoneally three times per week. PD0325901 (Selleck Chemicals, Cat. No. S1036) (1.5 mg/kg in 0.5% methylcellulose, 0.2% Tween80 in water) was administered daily by oral gavage. Treatments lasted 2 months, with endpoint MRI at 9 months. Durability scan was performed at 10 months with mice kept off treatment for 1-month.

### Statistical analysis

All in-vitro experiments were performed in at least three biological and three technical replicates. Statistical analysis and graphical representation of data were performed using GraphPad Prism software (version 10.4.1). Significance was determined by ordinary one-way ANOVA and Ordinary two-way ANOVA with Tukey’s multiple comparisons post-hoc test and with Unpaired t-tests. To compare the effect of BAY11-7082+PD0325901 combination treatment with PD0325901 single agent treatment on established mouse neurofibroma, a random-effects-model analysis on the log-transformed tumor volume data was performed as described previously (96). A *p*-value less than 0.05 was considered significant. Number of observations are reported in main text and Figure legends (mice, cells, samples).

### Study approval

All procedures were approved by the Institutional Animal Care and Use Committee at Cincinnati Children’s Hospital Medical Center.

## Data availability

All data are available in the main text or supplemental files. Data values for all graphs and reported means are provided in the Supporting Data Values file. RNA sequencing data is available through Gene Expression Omnibus (GEO) (GSE346553). High-resolution multiplexed CODEX fluorescence images can be interactively explored via the HALO Link Viewer (https://halo.cancer.gov/s/studies/19261). The full raw image TIFF files along with mapped single-cell segmentation masks and raw marker signal intensity values, are available from the corresponding author upon reasonable request.

## Acknowledgements

Supported by DOD-HT9425-1-0435 (to NR and JS), NIH R01 NS115438 (to DAL and NR), NIH R01 NS28840 (to NR), and a Children’s Tumor Foundation Young Investigator Award to RR. The authors would like to thank Katherine Chaney Scheffer for collecting primary cells and Dr. Jianqiang Wu for processing human tumor samples. Some work unrelated to the contents of this manuscript in NR’s lab is funded by Springworks, Boehringer-Ingelheim, and Gondola Bio. DAL is the co-founder and co-owner of biotechnology companies including NeoClone Biotechnologies, Inc., Discovery Genomics, Inc. (recently acquired by Immunsoft, Inc.), and B-MoGen Biotechnologies, Inc. (recently acquired by the Biotechne corporation). He is a co-founder of, and holds equity in, Luminary Therapeutics, Inc. He consults for Genentech, Inc., which has funded some of his research. The business of these companies is unrelated to the contents of this manuscript. This research was supported in part by the Intramural Research Program of the National Institutes of Health (NIH). The contributions of the NIH authors were made as part of their official duties as NIH federal employees, are in compliance with agency policy requirements, and are considered Works of the United States Government. However, the findings and conclusions presented in this paper are those of the authors and do not necessarily reflect the views of the NIH or the U.S. Department of Health and Human Services.

## Author contributions

RR, JS, DAL and NR conceived and designed the study and provided funding. RR, NK, EY, AV, JP, acquired data. RS provided human tissue. RR, EY, KC, ED, MK, SS analyzed and interpreted data. RR, EY, JP, DAL, JS, NR wrote, reviewed and/or revised the manuscript. NR supervised the study.

